# Bigger and Better? Representativeness of the Influenza A surveillance using one consolidated clinical microbiology laboratory data set as compared to the Belgian Sentinel Network of Laboratories

**DOI:** 10.1101/343236

**Authors:** Sigi Van den Wijngaert, Nathalie Bossuyt, Bridget Ferns, Laurent Busson, Gabriela Serrano, Magali Wautier, Isabelle Thomas, Matthew Byott, Yves Dupont, Eleni Nastouli, Marie Hallin, Zisis Kozlakidis, Olivier Vandenberg

**Affiliations:** Department of Microbiology, LHUB-ULB, Pole Hospitalier Universitaire de Bruxelles, Université Libre de Bruxelles, Brussels, Belgium; Sciensano, SD Epidemiology and Surveillance, Service ‘Epidemiology of infectious diseases’, Brussels, Belgium; Department of Clinical Virology, UCL Hospital NHS Foundation Trust, London, UK; NIHR UCLH/UCL Biomedical Research Centre, London, UK; Research Centre on Environmental and Occupational Health, School of Public Health, Université Libre de Bruxelles, Belgium; National Influenza Centre, Sciensano, Brussels, Belgium; UCL, Division of Infection and Immunity, Faculty of Medical Sciences, London, UK; Department of Population, Policy and Practice, UCL GOS Institute of Child Health, London, UK; The Farr Institute of Health Informatics Research, University College London, UK; Innovation and Business Development Unit, LHUB-ULB, Pole Hospitalier Universitaire de Bruxelles, Université Libre de Bruxelles, Brussels, Belgium

**Keywords:** Belgium, Influenza A, surveillance, clinical microbiology laboratory, sequencing

## Abstract

Infectious diseases remain a serious public health concern globally, while the need for reliable and representative surveillance systems remains as acute as ever. The public health surveillance of infectious diseases uses reported positive results from sentinel clinical laboratories or laboratory networks, to survey the presence of specific microbial agents known to constitute a threat to public health in a given population. This monitoring activity is commonly based on a representative fraction of the microbiology laboratories nationally reporting to a single central reference point. However in recent years a number of clinical microbiology laboratories (CML) have undergone a process of consolidation involving a shift towards laboratory amalgamation and closer real-time informational linkage. This report aims to investigate whether such merging activities might have a potential impact on infectious diseases surveillance. Influenza data was used from Belgian public health surveillance 2014-2017, to evaluate whether national infection trends could be estimated equally as effectively from only just one centralised CML serving the wider Brussels area (LHUB-ULB). The overall comparison reveals that there is a close correlation and representativeness of the LHUB-ULB data to the national and international data for the same time periods, both on epidemiological and molecular grounds. Notably, the effectiveness of the LHUB-ULB surveillance remains partially subject to local regional variations. These results illustrate that centralised CML-derived data are not only credible but also advantageous to use for future surveillance and prediction purposes, especially for automatic detection systems that might include multiple layers of information and timely implementation of control strategies.

## INTRODUCTION

Infectious diseases remain a serious public health concern in industrialized and low- and middle-income countries. Due to their considerable effect on global human demographics (1, 2) and the economy (3, 4), the public health community has developed many surveillance strategies and systems to improve infectious disease surveillance around the world. Even if diagnostic and computer resources have expanded considerably, infectious disease surveillance remains challenging. The 2009 H1N1 influenza pandemic and the recent Ebola outbreak in West Africa are few examples showing that infectious diseases cannot easily be predicted and modelled reliably in real-time (5, 6). Therefore the need for reliable and representative surveillance systems remains as acute as ever. One of the more established surveillance strategies, known as traditional public health surveillance of infectious diseases, is the use of reported positive results from sentinel clinical laboratories or laboratory networks to survey the presence of specific microbial agents known to constitute a threat to public health in a given population (7).

According to Colson et al., a total of 31 laboratory-based surveillance systems have been implemented in Europe for the purpose of global surveillance (8). Two such characteristic examples of surveillance systems implemented successfully using the above reporting strategy are the Health Protection Agency in England and Wales which counts infectious pathogens detected by hospital and specialist laboratories (9), and the surveillance systems of the Netherlands Reference Laboratory for Bacterial Meningitis (10).

In Belgium, such a strategy has been launched and implemented in 1983 by Sciensano, formerly known as Scientific Institute of Public Health (WIV-ISP), with the establishment of the sentinel laboratory network collecting information on the epidemiology of infectious diseases (11, 12). With 79-88 participating clinical microbiology laboratories (CMLs) across different geographical areas, the main objective of the Belgian Sentinel Network of Laboratories (BSNL) is to monitor the emergence and evolution of different infectious diseases over time, based on a representative fraction of the microbiology laboratories nationally (13, 14). With a ratio of 60% of participating laboratories to the total number of laboratories, the BSNL is able to describe trends and monitor changes in 12 groups of pathogens both at national and regional levels (15). However, in most European countries such laboratory surveillance is made on a voluntary base and is often not financially covered.

In parallel, the rationalisation of public health costs has led to the development of novel strategies for laboratories’ cost containment. In this perspective, a number of CMLs have undergone a process of consolidation involving a shift towards laboratory amalgamation and closer real-time informational linkage. Through this consolidation activity, an operational model has emerged with large centralized clinical laboratories performing on one central platform and one or several distal laboratories dealing locally only with urgent analyses. The increasing centralisation of diagnostic services over a large geographical region has given rise to the concept of “microbiology laboratories network” (16). The reduction in the number of small clinical laboratories and the aggregation of the remaining ones, may condition the ability to detect epidemiological changes. The sensitivity and representativeness of national surveillance systems should be therefore carefully monitored using coverage measures which indicate the proportion of the target population included within the surveillance system (15).

It is conceivable that the consolidated CMLs could become a cornerstone of public health models in the near future akin to the regional healthcare hospital networks with interactive surveillance for AMR control in France and cross-border regions (17). Due to the adoption of a 24/7 working scheme and improved automation, consolidated CMLs are also able to analyse a large influx of samples in the context of an outbreak investigation. In addition to the volume capacities and the large range of diagnostic tools, the ability of consolidated CMLs to access multiple different partners, geographies and clinical specialities can enhance their capabilities to provide advanced systems for disease surveillance and early recognition.

As there is the potential that such CMLs merging might have a consequence on infectious diseases surveillance, we used the availability of Influenza data to evaluate whether influenza infection trends could be estimated effectively from only one CML serving the Brussels area. The obtained data was compared to the available laboratory surveillance data provided by the directorate Epidemiology and public health of Sciensano in Belgium for the same time period.

## MATERIALS AND METHODS

### Location

The study took place in the Department of Microbiology of the Laboratoire Hospitalier Universitaire de Bruxelles - University Laboratory of Brussels (LHUB-ULB) serving 5 University Hospitals located on three geographical poles i.e. Centre, North and West of Brussels and representing close to 3,000 beds. Their catchment population covers a population of 700,000 inhabitants according to the Belgian statistical office, where the inhabitants of Brussels area are described for the year 2017 (18). Altogether, the LHUB-ULB performs annually more than 1,200,000 microbiology analysis including viral culture (12,000) and molecular diagnosis (26,000).

### Data collected

In our routine surveillance perspective, all patients diagnosed with flu infection by either ImmunoChromatographic Test (ICT), Lateral Flow Chromatography (LFC), molecular diagnostic tests or by cell culture methods are considered as notifiable cases of flu infection. These laboratory diagnosed cases of flu infection are transferred on a weekly basis to the BSNL by mail. The encoded variables include the diagnosed infectious disease, some patients demographic data allowing the identification of duplicates i.e. date of birth (or previously age), gender and postal code. In addition the specimen and its sample identification number, the diagnostic method and the date of diagnosis are recorded as well. For confidence reasons, all data transfers are kept anonymous towards the patient.

To assess the representativeness of the LHUB-ULB surveillance (for three seasons), LHUB-UB’s data collected from 1^st^ January 2014 to 31^st^ December 2017, were compared with all flu cases notification of the BSNL according to the geographical coverage and time distribution. In addition to BSNL epidemiological data’s, weekly reports on the incidence of clinical influenza-like illness (ILI) and virological data collected by sentinel general practitioners (SGPs) were also considered to better describe the epidemiological trends of flu at the regional and national level. Data from both sources were collected retrospectively and anonymised before analysis in a routine surveillance perspective. Ethics approval was granted by the Ethics Committee of the Saint-Pierre University Hospital. No written informed consent was collected.

To test the validity of LHUB-ULB data as a source for flu surveillance, the flu notification trends of the LHUB-ULB were compared with the trends of the overall BSLN network, BSLN network minus LHUB-ULB notification (BSLN-) and ILI consultation rate by Spearman’s rank correlation coefficient within each epidemic season and for the corresponding data over four years investigated at the national and regional level. All statistical tests were two-sided with the alpha set at 0.05. Analyses were done with the STATA/IC, version 14.

### Microbiology methods

To better assess the input of the LHUB-ULB as early warning lab for molecular surveillance, the genomes of 39 randomly selected Influenza A(H1N1)pdm09 isolates obtained from patients attending the emergency room of Saint-Pierre University Hospital, located in Brussels (01/02/2016 to 15/03/2016), were whole genome sequenced and compared to equivalent Influenza A(H1N1)pdm09 sequences deposited at the Epiflu database for the same location and period and against France, Germany, Netherland and UK-derived sequences from the same time period (Table of all isolate IDs used in this study are included in the Supplemental Digital Content 1).

All samples were initially screened for influenza A virus by reverse transcription-PCR targeting the matrix gene. A total of 39 influenza A virus-positive samples had their whole genome sequenced. RNA was amplified using a modified eight-segment method. Library preparations were generated as previously described (19) and short reads were assembled *de novo* (20). A neighbour joining phylogenetic tree was constructed using the Multiple Sequence Alignment Software v.7 (MAFFT) (21).

## RESULTS

From 2014 to 2017, 31,809 flu cases were declared by the sentinel laboratories to the BSNL. Among them, 3,186 infections were reported by the LHUB-ULB representing 38.75% (2,746/7,088) and 10.1% (3,186/31,809) of all influenza cases reported by the BSNL in the Brussels region and at the national level respectively. Figure 1 and 2 describe the incidence of flu infection’s notification by municipalities and the geographical representativeness of the LHUB-ULB in this notification for the year 2016 respectively. During the study period, the number of municipalities in which the LHUB-ULB represents at least 50% of coverage ranges from 21 (2017) to 32 (2016) representing 28.1% (21.6 to 36%) of the Belgium territory and 9.1% (8.1 to 10.4%) of the Belgium population (Supplemental Digital Content 2). Even if most of the municipalities are located in the Brussels region area; we note that each year the LHUB-ULB represents 100% of the notification of 5 to 10 municipalities located outside of the Brussels area either in Flanders or in Wallonia.

**Figure 1:**
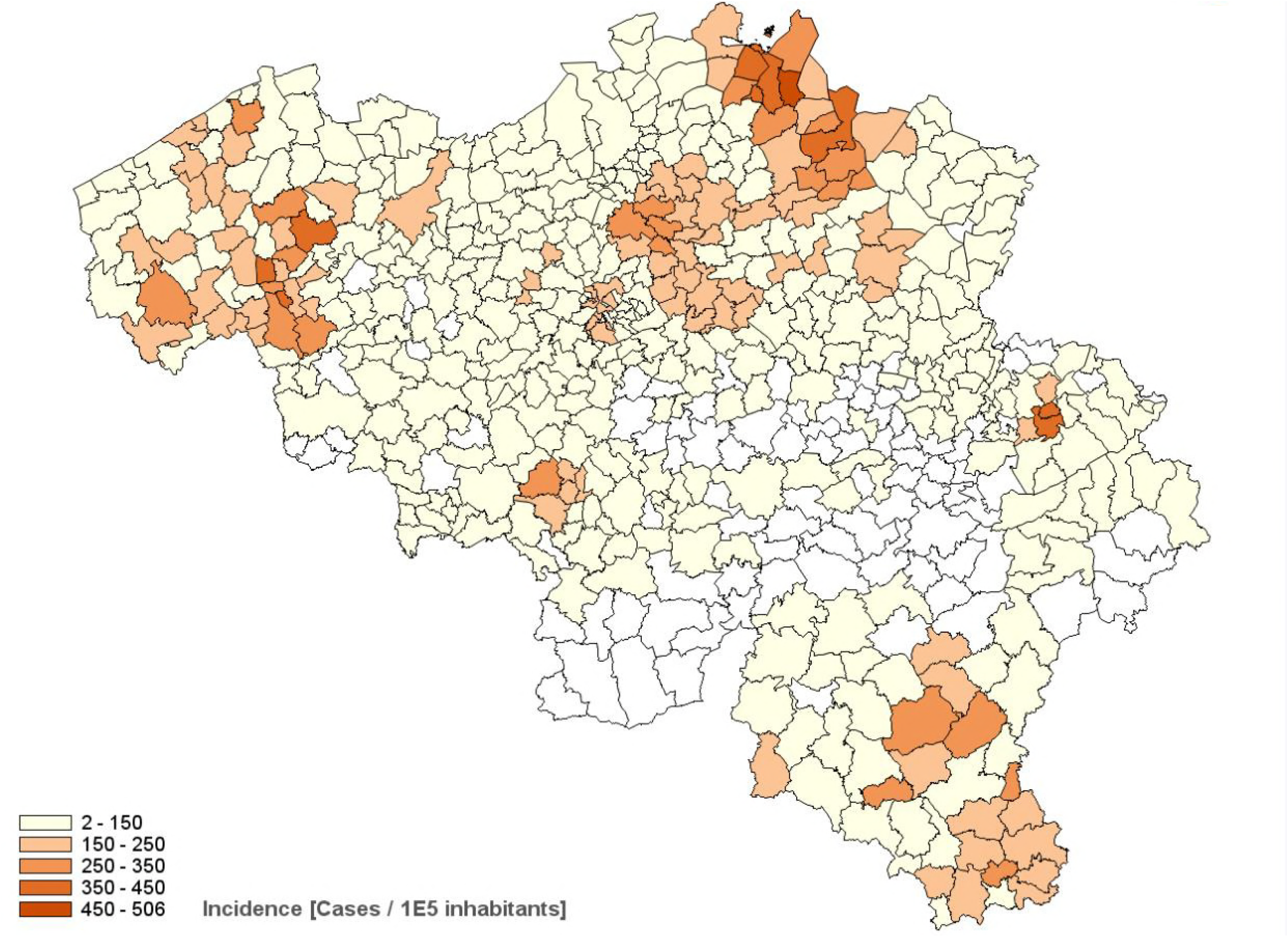
Incidence of the flu infection’s notification by municipalities in 2016

**Figure 2:**
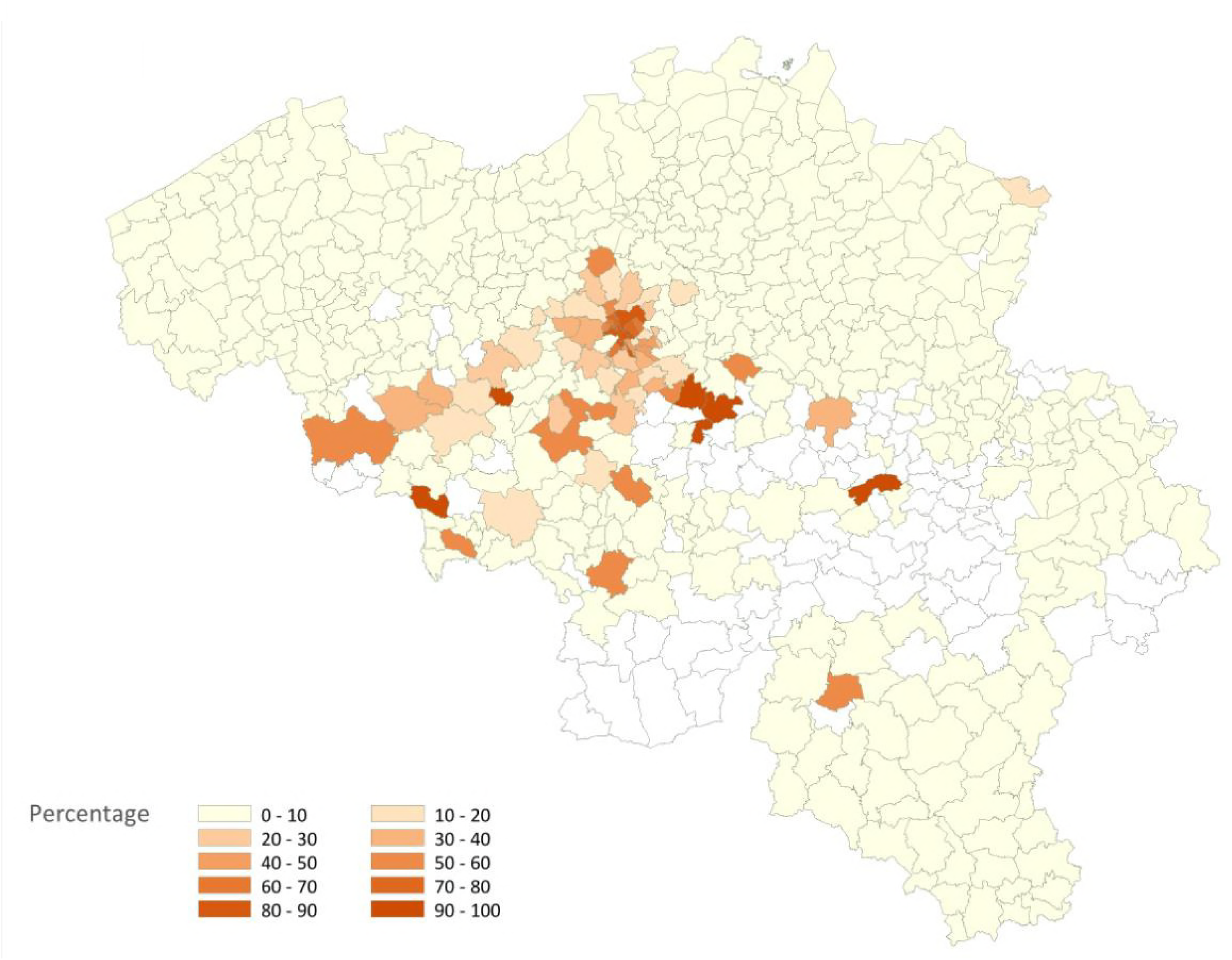
Representativeness of the LHUB-ULB data’s in the flu infection’s notification in the Belgian Sentinel Network of Laboratories in 2016

The representativeness of the LHUB-ULB to describe the trends of the flu notification at the regional and national level is described in figure 3. In order to better describe the flu trends, the incidence ILI based on data’s provided by the network of SGPs is also represented. Overall for the 4 years, the notifications provided by the BSLN closely follow the trends of the incidence of consultations by the SGP for ILI per 100,000 inhabitants with coinciding start, peak and end of the epidemics. In addition, the three sources of data show a comparable course over years and a high correlation within each season, i.e. Spearman’s rank correlation for each epidemic season ranged from 0.70 to 0.98 for the Brussels region and at the national level). However, LHUB-ULB’s data only were not able to reflect the trends of flu infection for the Flanders and Walloon regions, i.e. the lowest correlation (although still statistically significant) was observed in w40/14 to w39/15 for the Walloon Region (Supplemental Digital Content 3).

**Figure 3:**
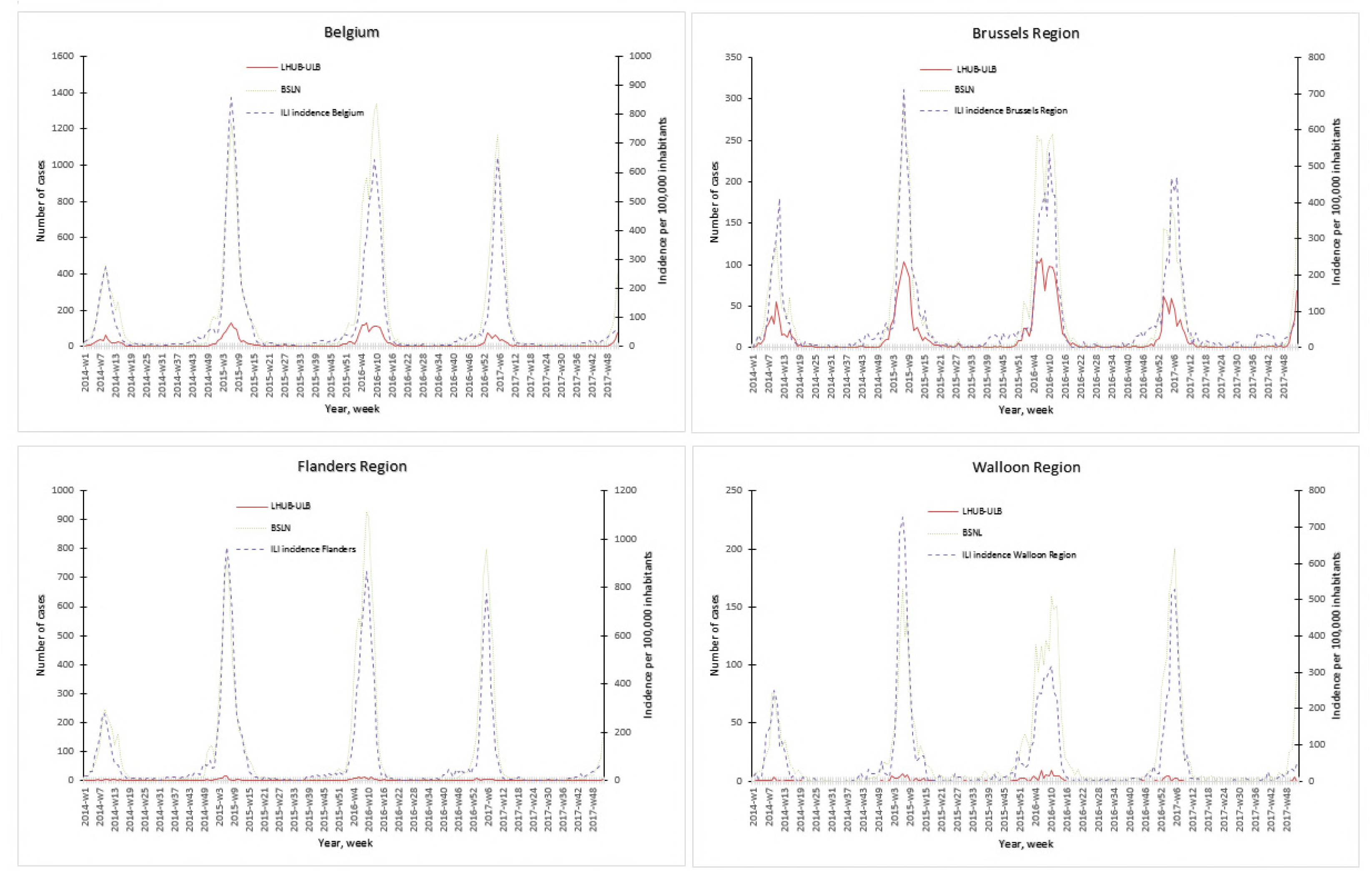
Representativeness of the flu surveillance in the Belgian Sentinel Network of Laboratories using one clinical microbiology laboratory data’s.

The representativeness of the LHUB-ULB at the microbiological level was also assessed by comparing the microbiological data’s with those of National Influenza Centre for the 3 consecutive flu seasons (2014-2017). Among the 2,309 respiratory samples sent by the network of SGPs during the study period and analysed at the National Influenza Centre (NIC), 1,197 (51.8%) were positive for influenza (70.0% influenza A, 23.1% influenza B). Thirty samples were co-infected with influenza A and B. Among the influenza A samples that were subtyped, 25.6% (236/922) were A(H1N1)pdm2009 and 70.4% (649/922) were A(H3N2). Thirty-six samples (7.6%) could not be subtyped due to their low viral load. Of the 277 influenza B samples analysed, 70.8% (196/277) belonged to the Victoria lineage and 28.8% (80/277) to the Yamagata lineage. During the same period, 72.5% (1,553/2,141) and 28.1% (601/2,141) of influenza viruses detected by the LHUB-ULB overall were type A and type B respectively. Difference in proportion with the NIC globally and over time were not significant (p>0.05) overtime and globally (Supplemental Digital Content 4).

The representativeness of the LHUB-ULB in terms of molecular epidemiology was assessed by the phylogenetic analysis of the sequenced isolates (Figure 4). The LHUB-ULB derived Hemagglutinin (HA) sequences (n=39) were compared to all sequences available from GISAID from Belgium for the same time period (n=17) and an equivalent number of sequences from the United Kingdom (n=56), France (n =126) and the Netherland (n =36). Most of the LHUB-ULB samples constitute distinct phylogenetic clusters, co-located with other samples with Belgium as a place of origin, within a background of seasonal Influenza A phylogeny. Preliminary results show that no significant genomic differences were observed within the LHUB-ULB samples or against deposited genomes from the same location and time period in Belgium, the Netherlands and France. However, most LHUB-ULB derived sequences form clusters distinct to the UK-derived samples. This observation would require larger sample numbers than the ones currently available and to be repeated for a number of seasons in order to be further validated. Figure 4b shows HA sequence analyses, though the LHUB-ULB samples did have the Neuraminidase (NA) genomic information available due to the whole genome sequencing, the availability of NA sequences in the public databases from the same samples was almost entirely unavailable.

**Figure 4:**
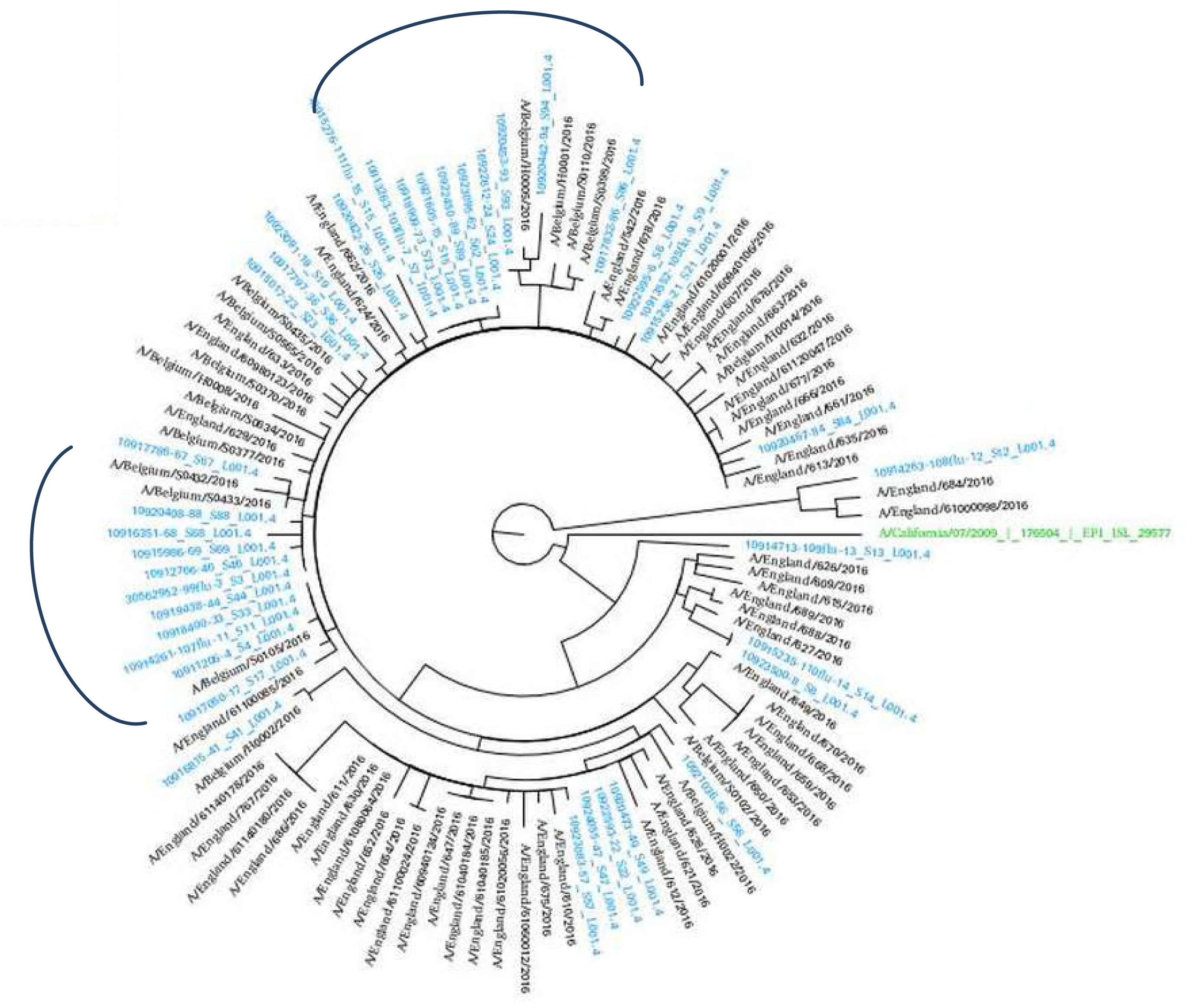
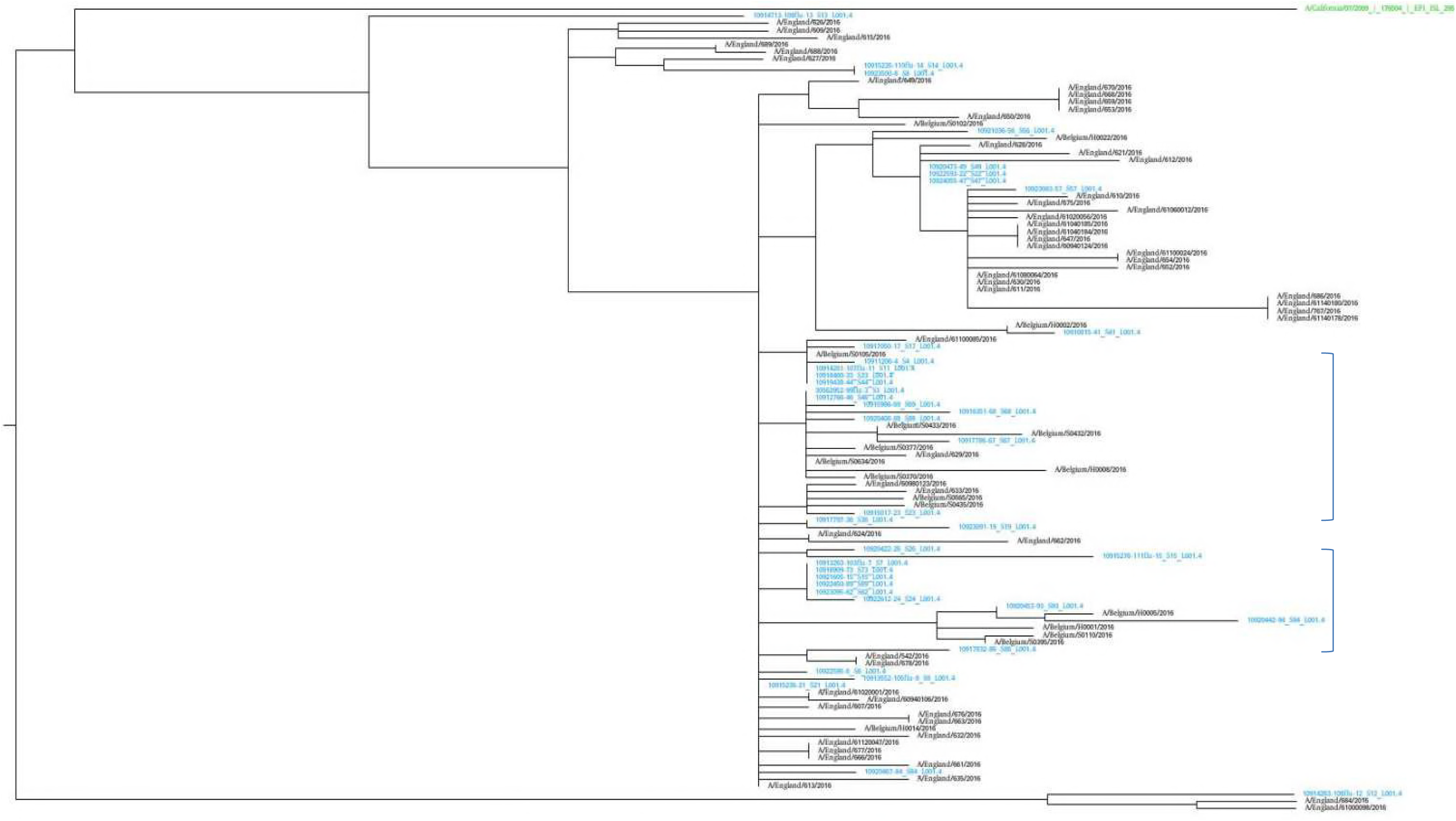
Phylogenetic trees of the HA gene sequences of the 39 Influenza A A(H1N1)pdm09 isolates from LHUB-ULB (shown in Blue), tested in this study compared to the reference strain (shown in green); all 17 sequences with Belgium as a place of origin for the same time period and equivalent number (56) of sequences deposited at the Epiflu database from the UK and same time period. The authors gratefully acknowledge the originating and submitting laboratories who contributed sequences used in the phylogenetic analysis to GISAID (http://www.gisaid.org). In both phylogenetic graphs (fig a and b), the two areas containing only samples derived from Belgium (including most of the LHUB-LUB samples) as place of origin are highlighted.

## DISCUSSION

In the 21st century, laboratory-based surveillance benefits from increased use of rapid diagnostic testing including ICT or LFT and multiplex PCR assay and increasingly rapid pathogen identification. The use of such advanced detection tools have dramatically cut the time to accurate diagnosis of infected patient but also increased the knowledge of epidemiological trends in comparison with older diagnostic methods based on culture and serology. The gradual integration of new detection and typing data into the European surveillance and alert systems represents one of the most exciting and challenging developments that could revolutionise the understanding of communicable diseases in the coming years (22).

However, such laboratory surveillance activity is often not financially covered and/or mandatory. Therefore, the cost containment leading to privatisation of laboratory medicine may have an impact on this laboratory activity which is not directly centred on patient care. For example, in Belgium, the consolidation process leads to a decrease of clinical laboratories from 496 in 1996 to 148 in 2017 with a shift toward large private laboratory structure. According to data on reimbursed microbiology tests obtained from the Belgian National Institute for Health and Disability Insurance (INAMI-RIZIV) for the period 2010–2015, on average 23.4% (range 22.5-24.3%) of all microbiological analysis (10,616,599 out of 45,355,005) performed were processed by private laboratories in regards to the 76.5% (range 77.4- 75.7 %) (34,696,723 out of 45,355,005) performed by university or hospital CMLs (Muyldermans G, unpublished data). The later being more traditionally involved in the non profit activities such as research and development and/or public health surveillance.

The use of data from large CMLs may file this gap that the privatization of the laboratory medicine may beget. In this study, the handling of the flu data from the LHUB-ULB reveals its attractive features that can facilitate an early detection of seasonal influenza epidemics. These results illustrate that data are not only credible but also advantageous to use for surveillance and prediction purposes, especially for an automatic detection system. Despite, the LHUB-ULB catchment area represents a small geographical area; its representativeness for the nation-wide data is striking. In the future, the extent of representation will be further improved when data are collected from more consolidated laboratories.

In addition, this study confirms the lack of covering by BSLN in some municipalities located in Flanders or in Wallonia (15). In our study, the LHUB-ULB represented each year 100% of the notification in several municipalities located in these regions. The use of laboratory information’s provided by one consolidated CML thanks to new molecular tools show to be a good complement of the information’s provided by the SGPs and Hospital network and use by the NRC influenza (23, 24). The real-time integration of consolidated CMLs into the public health surveillance system would help the monitoring of influenza activity (intensity, duration, severity,…) all over the year, the determination of type and subtypes of circulating strains and their antigenic and genetic characterization. A such integration could also to contribute to the annual determination of the influenza vaccine content, the monitoring of resistance to antivirals and the detection of new potentially pathogenic influenza viruses (25).

In this frame, the use of high-throughput whole genome sequencing platforms available in large CMLs network such as met in the LHUB-ULB demonstrates its potential for molecular epidemiological surveillance. Because, early detection of epidemics is a key element to prevent loss of (quality of) life and its economic and material impact, such molecular surveillance would gain in efficiency through automated real-time monitoring and reporting to public health authorities from the regional to the European levels. Other European countries have started demonstrating a clinical benefit from such data collected initiatives, as is the case in the UK at the National Mycobacterial Reference Service in Birmingham (26) and at UCLH with the integration of near real-time, whole genome sequencing utilised for the purposes of HIV and Influenza surveillance (19, 27).

However the availability of new microbial typing and detection techniques and culture-independent diagnostic methods, brings about a fundamental change in the way data has to be handled. These approaches are high-throughput and data-rich and create systematic stresses in the collection, analyses and safe handling of the generated data (28). For example one current obstacle towards clinical translation is that most algorithms in use need some programming expertise, together with specialized servers to handle and store all of the data. The generation of user-friendly informatics tools to effectively analyse high-throughput genomic data will be essential to the successful clinical application of genomic technology. In addition there are systemic aspects that also need to be addressed, such as the number of additional molecular or genomic testing parameters, regardless of the method, which can be supported routinely by the existing electronic health records that provide the architectural framework. These aspects can include the ordering of the test, the receiving of a document that summarizes the clinical interpretation, and storage of the interpretation (29). The integration of molecular genetic data to clinical and/or epidemiological data creates a challenge that requires interactive, information-sharing workspaces rather than uni-directional centralised reporting, such as those deployed in the TYPENED approach (30).

## CONCLUSION

Our results suggest that consolidated CMLs represent a wealth of information, including data usable for public health surveillance. The advent of real-time sequencing of organisms, their direct integration in surveillance tools at the regional, national and European levels would lead to real-time detection and alert, allowing the rapid prioritization of public health threats and the timely implementation of control strategies

## ACKNOWLEDGMENTS

We wish to thank the personnel of the Department of Microbiology of LHUB-ULB for its daily technical assistance. This work has been supported in part by a UCL Global Engagement Fund award to Prof Vandenberg and Dr Kozlakidis, which is gratefully acknowledged. The authors would like to thank Gaetan Muyldermans for his help in the preparation of the manuscript. This work was partly presented at the 28th European Congress of Clinical Microbiology and Infectious Diseases. 2018 April 21-24. Madrid, Spain. p.112: P0001. No conflict of interest to declare.

